## Supplementary Text for "A neural-mechanistic hybrid approach improving the predictive power of genome-scale metabolic models"

### Supplementary Information

**Wt-solver equations** **2**
Figure S1. Computing steady-state fluxes with the Wt-solver 4
**AMN-Wt architecture** **5**
Figure S2. Different weights for different uptake fluxes provided by exact bounds (EBs) 5
Figure S3. Branch point metabolite flux ratio 6
**LP-solver equations** **8**
Figure S4. Matrices used with the LP (EB) method 11
Figure S5. Matrices used with the LP (UB) method 12
**QP-solver equations** **13**
**MM solvers benchmarking** **16**
Figure S6. MM solvers architectures and performances 16
**AMNs benchmarking varying hyperparameters** **17**
Figure S7. Hyperparameters for AMN's neural Layer 17
**AMNs benchmarking with independent test sets and additional metabolic models** **18**
Table S1. Benchmarking MMs, ANNs and AMNs 18
**AMNs benchmarking with gene knockouts and multiple measured fluxes** **21**
Figure S8. AMN performance on multiple fluxes dataset 22
**AMN-Reservoir prediction performance** **23**
Figure S9. Performances of AMN-Reservoir using predicted $V_{in}$ as input to FBA 23
**ANN training set sizes** **24**
Figure S10. Loss and regression coefficient for training sets of increasing sizes 24
**Experimental workflow** **25**
Figure S11. Experimental workflow pipeline 25
**Terminology** **26**
Table S2. Vectors and matrices notations used in figures and equations 26

**Supplementary references** **28**

**Wt-solver equations**

The Wt-solver equations, inspired by the work of Nilsson *et al.*[1] for signaling modeling, are detailed below for the specific example given in Figure S1.

With the usual FBA method (Figure S1.a), we obtain the flux distribution maximizing the flux of the $v_3$ reaction (representing a classical 'biomass' reaction) with an uptake reaction $v_1$ by solving a linear program (Figure S1b).

In the Wt-solver method, we first translate the model into a neural-network-like architecture (Figure S1c). Precisely, in Figure S1c we start from an initial set of given fluxes ($v_1$ = 0.1) and then propagate knowledge about the fluxes through the entire network, each layer corresponding to one step in a discrete flux propagation. Mathematically, each layer is composed of two simple operations that update the *M* and *V* vectors, respectively representing metabolites production rates and reaction fluxes. Those operations are repeated until convergence. The network shown in Figure S1c is the unrolled representation of an RNN-like network depicted in Figure S1d. The matrices used in the Wt-solver are given in Figure S1e. The weight matrix $W_r$ can be computed from the steady state flux values if they are known, or learned through training from provided reference fluxes (*cf.* section 'AMN-Wt architecture'). For instance, taking the example of Figure S1, the weight matrix $W_{\mathrm{r}}$ can be either computed from steady state fluxes, computed by FBA (*cf*. legend of Figure S1f), or learned after setting $v_1$ *= 0.1* and searching weights for which $v_3$ *= 0.5*. We note, the steady state fluxes after iterating 30 times (or more, here we stopped at 30 because the reference data values were reached) are equal to those obtained in the reference data (*cf*. Figure S1 panels b and f).

**a**

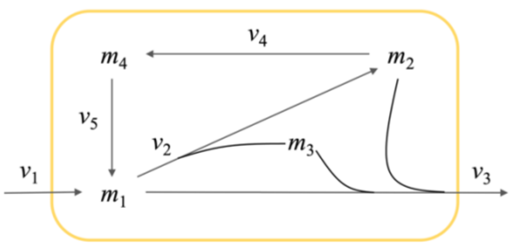


**d**

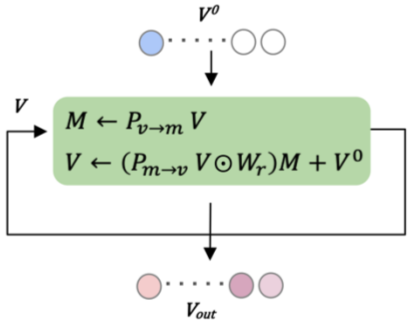


**b**

**Solve with Linear Programming:**

$$\begin{aligned} &Max(c^T V) \\ &\text{Subject to:} \\ &\quad SV = 0 \\ &\quad 0 \le V \le V_{in} \end{aligned}$$

$$\text{where: } V_{in} = (0.1, -, -, -, -)^T, \quad V = (v_1, \cdots, v_5)^T, \quad c = (0, 0, 1, 0, 0)^T$$

$$\text{and } S = \begin{pmatrix} 1 & -1 & -0.13 & 0 & 1 \\ 0 & 1 & -0.07 & -1 & 0 \\ 0 & 1 & -0.36 & 0 & 0 \\ 0 & 0 & 0 & 1 & -1 \end{pmatrix}$$

(columns $v_1, v_2, v_3, v_4, v_5$; rows $m_1, m_2, m_3, m_4$)

**Steady state solution:**

$$V_{out} = (0.1, 0.18, 0.5, 0.15, 0.15)^T$$

**e**

$$P_{v\to m} = ReLU(S) = \begin{pmatrix} 1 & 0 & 0 & 0 & 1 \\ 0 & 1 & 0 & 0 & 0 \\ 0 & 1 & 0 & 0 & 0 \\ 0 & 0 & 0 & 1 & 0 \end{pmatrix}$$

$$P_{m\to v} = ReLU\left(\left[\frac{-1}{z_i s_{j,i}}\right]\right) = \begin{pmatrix} 0 & 0 & 0 & 0 \\ 1 & 0 & 0 & 0 \\ \frac{1}{0.39} & \frac{1}{0.21} & \frac{1}{1.08} & 0 \\ 0 & 1 & 0 & 0 \\ 0 & 0 & 0 & 1 \end{pmatrix}$$

**f**

$$W_r = \begin{pmatrix} 0 & 0 & 0 & 0 \\ 0.74 & 0 & 0 & 0 \\ 0.26 & 0.20 & 1 & 0 \\ 0 & 0.80 & 0 & 0 \\ 0 & 0 & 0 & 1 \end{pmatrix}$$

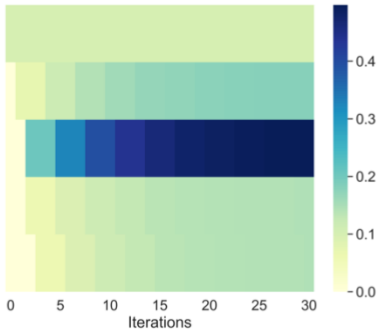


$$V^{30} = (0.1, 0.18, 0.5, 0.15, 0.15)^T$$

**c**

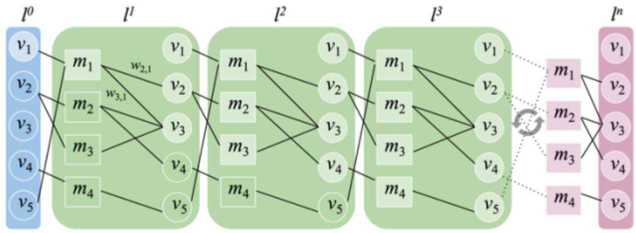

#### Figure S1. Computing steady-state fluxes with the Wt-solver

***a.*** *Simple toy stoichiometric model. The model is unidirectional, all flux values are positive. $v_1$ represents a nutrient uptake flux and $v_3$ the objective (e.g. the biomass flux). This toy network was inspired from the upper glycolysis pathway found in iML1515*[2]*, downloaded from BiGG*[3]*.* ***b.*** *Steady-state solution fluxes maximizing $v_3$. At steady state, the reaction fluxes ($v_i$) must satisfy stationarity conditions that guarantee mass balance of all metabolites, this is depicted by the equation $SV = 0$, where $S$ is the stoichiometric matrix representing the connectivity of the model and $V$ the vector of fluxes to be calculated. $V_{in}$ is the medium represented by a vector of nutrient uptake fluxes (here $v_1$= 0.1, symbol "—" indicates that no value is provided, in practice one uses an 'infinity' value to represent an unbounded flux). The steady-state solution $V_{out}$ is calculated by solving a linear program maximizing the objective $c^T V = v_3$, here the Cobrapy package was used to compute $V_{out}$, by making use of a Simplex-solver algorithm.* ***c.*** *Unrolled neural network built from the stoichiometric model. In the initial layer ($l^0$) only $v_1$ has a value. In layer 1, $v_1$ value is passed to $m_1$, the production flux for metabolite $m_1$. Subsequently a fraction ($w_{21}$) of $m_1$ goes to $v_2$ and the other fraction ($w_{31}$) to $v_3$. In layer 2, $v_1$ continues to feed $m_1, v_2$ is passed on to $m_2$ and $m_3$ , and then goes to $v_3$ and $v_4$. In layer 3, $m_4$ receives input from $v_4$ which in turn activates $v_5$. The unrolling is iterated until the values for the metabolites production rates and reaction fluxes converge.* ***d.*** *Recurrent neural network (RNN) representation. Vectors $M$ and $V$ respectively represent metabolites production rates and reaction fluxes. At each iteration step, $M$ and $V$ are computed using matrices $P_{v \to m}$ and $P_{m \to v}$ of panel e. When a metabolite is the substrate of several reactions (like $m_1$ and $m_2$), each reaction gets a fraction of the metabolite production flux, this is depicted in matrix $W_r$ (r indicates this matrix is used in recurrence). The matrix $W_r$ can be computed from the steady state fluxes of all reactions or learned through training. The operator $\odot$ stands for element-wise matrix product (Hadamard product).* ***E.*** *Neural network matrices. $P_{v \to m}$ is the matrix to compute metabolite production fluxes from reaction fluxes, $P_{m \to v}$ is a matrix to compute reaction fluxes from the production rates of the reaction substrates. $ReLU(x) = max\ (0, x)$, $s_{j,I}$ corresponds to the value of S at the $j^{th}$ row (metabolite) and $i^{th}$ column (flux) and $z_i$ is the number of strictly negative elements in column i of S.* ***f.****: Providing the FBA steady state solution, the weight matrix is computed as follows: $w_{21} = v_2/(v_1+v_5)$, $w_{31} = v_3/(v_1+v_5)$, $w_{23} = v_3/v_2$ and $w_{24} = v_4/v_2$. Heatmap obtained for n=30 iterations of the RNN of panel d running with the toy model of panel a.*

#### AMN-Wt architecture

As shown in Figure 1 in the main paper, the AMN-Wt architecture (like AMN-LP or AMN-QP) takes as input a vector ($V_{in}$) representing exact bounds (EBs) or upper bounds (UBs) for uptake fluxes. In both cases an initial vector $V^0$ is computed via the weight matrix $W_i$ ($V^0 = W_i V_{in}$). We recall from the previous section that AMN-Wt also comprises a weight matrix $W_r$ which can be learned through training. In the EB case, $W_i$ is not trainable and is just a mapping of $V_{in}$ into $V$. Consequently, in this case only the matrix $W_r$ is learned during training. In the UB case, the weights in $W_i$ are trainable and transform the upper bounds $V_{in}$ into exact bounds in $V^0$. Consequently, with UB, both $W_i$ and $W_r$ are learned during training. It turns out with EB that a single set of weights (matrix $W_r$) cannot handle all the elements of a training set when the network contains internal reaction fluxes depending on at least two metabolite uptake fluxes. Figure S2 below shows such an example. Consequently, AMN-Wt cannot be used to process EB training sets.

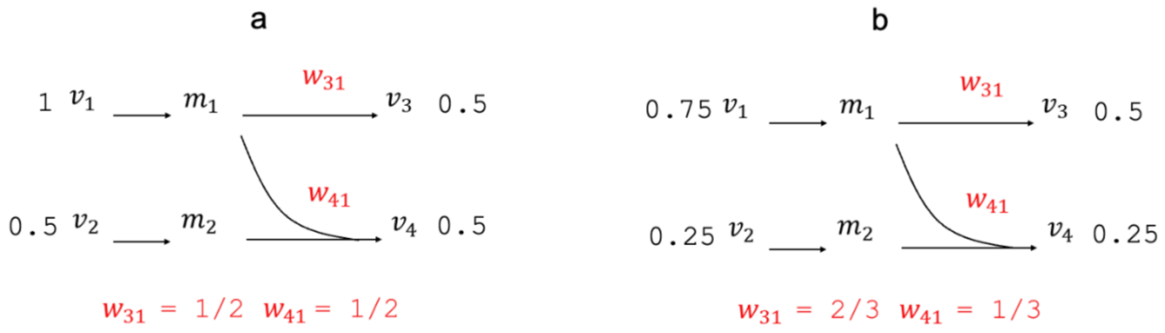


**Figure S2. Different weights for different uptake fluxes provided by exact bounds (EBs)**
*In the two cases all flux values ($v_i$) satisfy the steady state constraints (SV = 0, cf. Figure 1b). Following the equations provided in Figure 1d, the production rates for $m_1$ and $m_2$ are respectively 1 and 0.5 in (**a**) and 0.75 and 0.25 in (**b**). The reaction for flux $v_4$ is taking two substrates $m_1$ and $m_2$ and the value for $v_4$ is the minimum metabolite production rate (i.e., the rate limiting among $m_1$ and $m_2$). Consequently, the value for $v_4$ is 0.5 (**a**) and 0.25 (**b**). Therefore, the fraction ($w_{41}$) of $m_1$ contributing to $v_4$ is 1/2 (**a**) and 1/3 (**b**). The weights are different in panel a and b as they depend on the uptake flux values.*

In the UB case, the uptake flux upper bounds are first transformed into exact bounds with the matrix $W_i$ learned during training. In this instance, a vector $V^0$ is calculated via $W_i$ for each element of a training set, and as observed in Table S1, large training sets can be processed with a single set of weight $W_r$. Returning to the example of Figure S2, assuming we measure fluxes $v_3$ and $v_4$, then for any non-null weights $w_{31}$ and $w_{41}$, it is easy to find exact bounds for $v_1$ and $v_2$: $v_1 = v_3 / w_{31}$ and $v_2 = 2v_4 - v_3 w_{41} / w_{31}$. More generally, as shown in Table S1, when running AMN-Wt for many training sets with upper bounds for uptake fluxes for both *E. coli* core[4] and iML1515[2] models, we always find solutions with losses around 0.001 and for which the regression coefficients between AMN predicted growth rates and Simplex calculated growth rates are above 0.98.

Remains the question of whether or not a single set of weights is realistic from a metabolic kinetics point of view. Weights arise at branching points where metabolite fluxes can contribute to two or more reactions. For instance, taking the example of Figure S2, the metabolite production flux $m_1$ is spliced into $w_{21}m_1$ and $w_{31}m_1$. The question we therefore have to answer is: given different production rates for branch point metabolites, are the weights conserved?

Without lack of generality and for simplicity, we consider below a branch point metabolite with different production fluxes contributing to two reactions. Let $V$ be the production flux of that metabolite, and let $V_1$ and $V_2$ be the fluxes of the two reactions, we necessarily have $V = V_1 + V_2$. Now according to Michaelis-Menten equations:

$$V_1 = \frac{V_{max,1}\, S}{S + K_{m,1}},\ V_2 = \frac{V_{max,2}\, S}{S + K_{m,2}} \tag{S1}$$

where $V_{max,1} = E_1 k_{cat,1}$ ($V_{max,2} = E_2 k_{cat,2}$), $E_1$ ($E_2$) being the concentration of the enzyme catalyzing the reaction, $S$ the concentration of the branch-point metabolite, and $k_{cat,1}$ ($k_{cat,2}$) and $K_{m,1}$ ($K_{m,2}$) the turnover rate and the Michaelis constant of the reaction. Using these notations, we have:

$$V = \frac{V_{max,1}\, S}{S + K_{m,1}} + \frac{V_{max,2}\, S}{S + K_{m,2}} \tag{S2}$$

We note that $S$ can be computed from $V$ by solving the quadratic eq. S2. As shown in Figure S3 below, a numerical simulation for different values of the kinetics parameters shows that the ratio $(V_1 / V)$ remains constant (slope of Figure S3 equals 1) even when the production flux $V$ is changed by several orders of magnitude. Consequently, a single set of weights can fit a training set as long as the kinetics parameters $V_{max,1}$ ($V_{max,2}$) and $K_{m,1}$ ($K_{m,2}$) do not change with the production flux $V$.

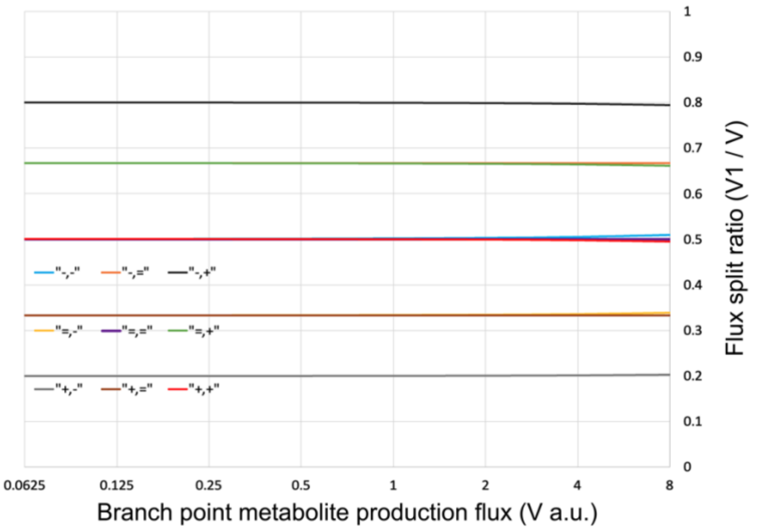


**Figure S3. Branch point metabolite flux ratio**

*Here a branch point metabolite contributes to two reactions with fluxes $V_1$ and $V_2$. The kinetics parameters for reaction $V_1$ are arbitrarily set to $V_{max1}$ = 1000 (a.u) and $K_{m1}$ = 100 (a.u). The values for $V_{max,2}$ and $K_{m,2}$ are those of $V_{max,1}$ and $K_{m,1}$ halved, equal or doubled. Nine cases are considered from $V_{max,2}$ = ½ $V_{max,1}$, $K_{m,2}$ = ½ $K_{m,1}$ ("-,-") to $V_{max,2}$ = 2 $V_{max,1}$, $K_{m,2}$ = 2 $K_{m,1}$ ("+,+").*

According to Figure S3, flux split ratios are conserved for nutrients leading to different metabolite production fluxes if the Michaelis-Menten kinetics parameters ($V_{max}$ and $K_m$) of the enzymes catalyzing the reactions involved in the split remain constant. However, Chubukov *et al*.[5] have shown experimentally that it was not the case (for *B. subtilis*) and different nutrients do provide different ratios. This behavior is due to varying enzyme activities, which themselves depend on enzyme concentrations, post-translational modification, and gene regulations. Chubukov *et al*.[5] showed with experimental evidence that different nutrients give rise to different concentrations for many enzymes, implying that nutrients do have an effect on gene regulations. We note that even though weights in $W_r$ do not have a physical meaning, AMN-Wt still exhibits excellent performances, showing that the consensual $W_r$ matrix and the initial $V^0$ vector (computed through a neural layer from the upper bound $V_{in}$) are performant enough. We also note that the weight issue does not arise with AMN-LP and AMN-QP as these architectures do not rely on flux split ratios.

**LP-solver equations**

We recall that the LP-solver makes use of Hopfield-like networks, which is a long-standing field of research[6] inspired by the pioneering work of Hopfield and Tank[7]. Later, these Hopfield-like networks were showcased to perform well for solving linear programs[8] and simpler and more efficient solutions were developed over the years[9,10]. It is important to point out at this stage that these Hopfield-like networks are non-trainable networks and differ from classical neural networks used in ML. The Hopfield-like networks are instead recurrent procedures iteratively updating the solution of linear programs.

The constrained linear optimization problems EB and UB are specific cases of the general problem described in Yang *et al.*[9] which can be written as:

$$min: c^T x \quad \text{(S3)}$$

s.t. $Ax = b$

$Bx \leq d$

where *x* is the vector of unknown (size *n*) to be calculated, *c* is the objective vector (also of size *n*), *A* is a (*m* x *n*) matrix of rank *m* (and therefore non-null) and *B* a *m x n* matrix. Translated to FBA problems, *x* is the flux vector (*V*), *c* the vector corresponding to the objective function, *A* is related to the stoichiometric matrix, *B* is a matrix that extracts exchange fluxes from the full flux vector and *d* is a vector of constraints on exchange fluxes.

Consequently, to solve a FBA problem *A, B, c, d, b* and *x* take the following values in the EB case:

$$A = S_{int}, B = -I_n, b = -b_{FBA}, d = 0\,, c = -c_{FBA}, x = V$$

According to Yang *et al.*[3] , gradients for V and its dual U in EB case can be written as:

$$\nabla V = (I_n - P)\left[\, c_{FBA} - {S_{int}}^T R\right] + QV \quad \text{(S4)}$$

$$\nabla U = \frac{1}{2}(U \; - R)$$

where:

$$R = ReLU(U + S_{int}V + b_{FBA})$$

$$Q = S_{int}^T\left(S_{int}S_{int}^T\right)^{-1}$$

$$P = QS_{int}$$

As $S_{int}S_{int}^T$ has to be invertible, it is important that $rank(S_{int}) = (S_{int})$ . To ensure this point, $S_{int}$ was converted to its row echelon form and rows with null values were removed. Consequently, prior to any computation of the LP solver, one has to preprocess the matrices in row echelon form.

The linear problem in the case where uptake fluxes values are unknown (UB method) can be written as:

$$max: c^T{}_{FBA}V \tag{S5}$$

s.t. $S_{int,In}V \geq -b_{FBA,0}$

$SV = 0$

The matrix $S_{int,In}$ is obtained by concatenation of $S_{int}$ and $-I_n$ (see Figure S5c). This matrix ensures the respect of two inequalities; entry fluxes are inferior to the upper bound, fluxes are positive. Consequently, $b_{FBA,0}$, is the concatenation of the upper bounds vector $b_{FBA}$ and a zero vector of size *n.*

The dual form of eq. S5 being:

$$min: -b^T{}_{FBA,0}U \tag{S6}$$

$st$: $S^T{}_{int,In}U \leq c_{FBA}$

$U \leq 0$

Thus, *A, B, c, d, b* and *x* take the following values: $A = S$, $B = -S_{int,In}$, $b = 0$, $d = b_{FBA,0}$ , $c = -c_{FBA}$, $x = V$

We note $c = -c_{FBA}$ since $\min\ (-c_{FBA}V)$ is equivalent to $\max\ (c_{FBA}V)$. The matrices A, B and vectors *b, d* take different forms depending if we are in the EB or UB cases. More details can be found in Figure S4 and Figure S5.

Similarly, to the EB case, gradients for U and V are:

$$\nabla V = (I_n - P)[\, c_{FBA} - S^T R] + Q\big(S_{int,In}V + b_{FBA,0}\big) \tag{S7}$$

$$\nabla U = \tfrac{1}{2}(U\ - R)$$

where:

$$R = ReLU(U + SV)$$

$$Q = S^T(SS^T)^{-1}$$

$$P = QS$$

Yang *et al.*[9] proved in their paper that *x*, the variable of the primal problem, and *y* ,the variable of the dual problem, (*V* and *U* using FBA notations) can be calculated iteratively starting with arbitrary values for $V^{(0)}$ and $U^{(0)}$:

$$V^{(t+1)} = V^{(t)} - dt\ \nabla V \tag{S8}$$

$$U^{(t+1)} = U^{(t)}\ -\ dt\ \nabla U$$

where $t$ is the iteration number, $dt$ the learning rate, and the derivatives are equation (S4) for EB and (S7) for UB.

The equivalence between (S3) and (S8) is proved by the first Lemma of Yang *et al.*[9] which states that $V^*$ is a solution of (S3) if and only if there exist $U^*$ such that:

$$(I - P)(-c - B^T\, U^*) - Q(A\, V^* - b) = 0 \quad \text{(S9)}$$

$$ReLU(U^* + B\, V^* - d) - U^* = 0$$

with $I$ the identity matrix of adequate size for $P$. According to the 2nd Theorem in Yang *et al.*[9], any initialization will converge to an equilibrium point. This Theorem also states that the convergence trajectories are asymptotically stable if there is a unique equilibrium point. In practice it is almost never the case with metabolic networks because the optimum is rarely unique.

Note that the work from Yang *et al.*[9] allows quadratic optimization, thus this method could be used with a fitting term similar to the one described in the QP method.

a

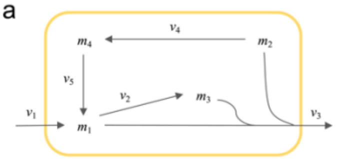


$$S = \begin{array}{c} \\ \end{array}\begin{matrix} v_1 & v_2 & v_3 & v_4 & v_5 \end{matrix}$$

$$S = \begin{pmatrix} 1 & -1 & -0.13 & 0 & 1 \\ 0 & 1 & -0.07 & -1 & 0 \\ 0 & 1 & -0.36 & 0 & 0 \\ 0 & 0 & 0 & 1 & -1 \end{pmatrix} \begin{matrix} m_1 \\ m_2 \\ m_3 \\ m_4 \end{matrix}$$

$$V^T = (v_1, v_2, v_3, v_4, v_5)$$

$$b_{FBA}^T = (0.1, 0, 0, 0)$$

$$Max: c_{FBA}^T V$$

$$s.t.\ S_{int} V = -b_{FBA}$$

$$-V \leq 0$$

$$\nabla V = (I_n - P)[c_{FBA} - S_{int}^T R] + QV$$

$$\nabla U = \frac{1}{2}(U - R)$$

b

Ensure that $S_{int}V$ = -$b_{FBA}$

$S_{int}$ is used for equality

All intake fluxes are turned to zero in S

$$\begin{matrix} v_1 & v_2 & v_3 & v_4 & v_5 \end{matrix}$$

$$S_{int} = \begin{pmatrix} 0 & -1 & -0.13 & 0 & 1 \\ 0 & 1 & -0.07 & -1 & 0 \\ 0 & 1 & -0.36 & 0 & 0 \\ 0 & 0 & 0 & 1 & -1 \end{pmatrix} \begin{matrix} m_1 \\ m_2 \\ m_3 \\ m_4 \end{matrix}$$

Ensure that -V ≤ 0

-$I_n$ is used to ensure fluxes positivity

c

$$R = \begin{cases} ReLU(U + S_{int}V + b_{FBA}) & if\,EB \\ ReLU(U + SV) & if\,UB \end{cases}$$

$$Q = \begin{cases} S_{int}^T (S_{int} S_{int}^T)^{-1} & if\,EB \\ S^T (SS^T)^{-1} & if\,UB \end{cases}$$

$$P = \begin{cases} QS_{int} & if\,EB \\ QS & if\,UB \end{cases}$$

**Figure S4. Matrices used with the LP (EB) method**

***a.*** *We show here an example for the same model as in Figure S1 in the scenario of known uptake fluxes (EB). All the information about uptake fluxes is contained in $b_{FBA}$. P, Q and R are defined in panel c (in this case using the formulations for exact bounds, EB). ∇V and ∇U are gradients respectively for the fluxes and metabolites shadow prices (as in the eq. S4)* ***b.*** *In the EB case, the only inequality constraint to verify is the positivity of fluxes (highlighted in red), so we use -$I_n$, the identity matrix of size n (number of fluxes), multiplied by -1, as the matrix B in Yang et al. formulation. To verify the equality constraints, coefficients of $v_1$ are zeroed out in $S_{int}$ because only the input flux $v_1$ of the metabolite $m_1$ is known, thus the matrix A in Yang et al. formulation is $S_{int}$ in our formulation, and $S_{int}V$ = -$b_{FBA}$ ensures the respect of equality constraints.* ***c.*** *Reminder of P, Q and R when using exact (EB) or upper bounds (UB).*

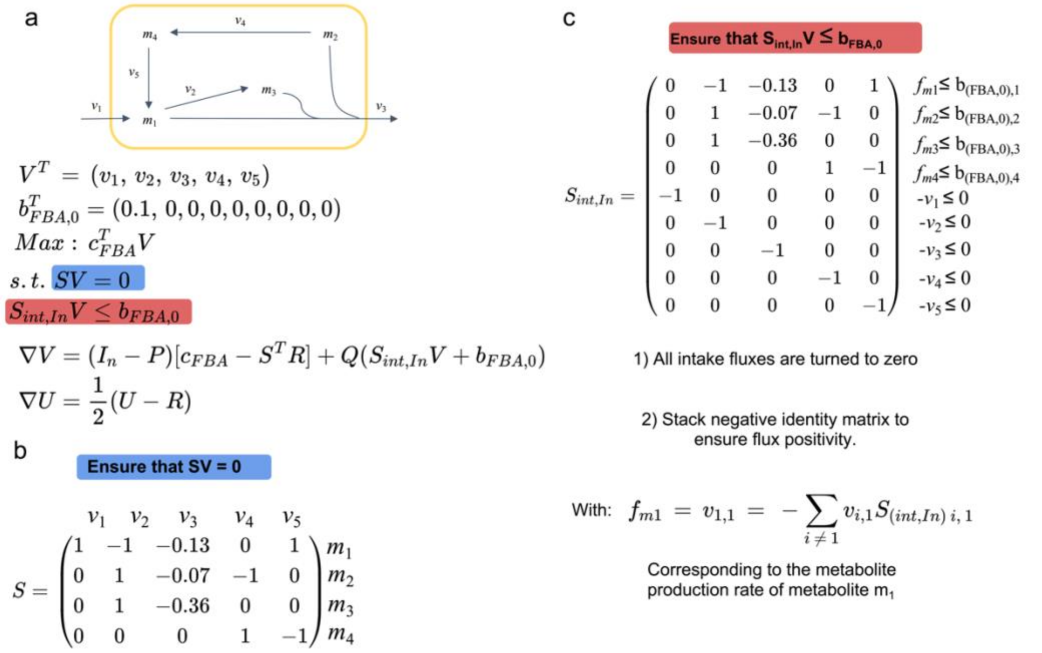


**Figure S5. Matrices used with the LP (UB) method**

***a.*** *The example is the same as in Figure S1 and Figure S4. In this scenario, uptake fluxes are unknown (UB), consequently upper bounds are contained in $b_{FBA,0}$ and their values are fixed arbitrarily. Note that, in $b_{FBA,0}$ , an upper bound is different from zero if and only if the corresponding metabolite is in the medium (only $m_1$ here). P, Q and R are shown in. Figure S4, panel c (in this case using the formulations for upper bounds, UB). $\nabla V$ and $\nabla U$ are gradients respectively for the fluxes and metabolites shadow prices (as in the eq. S7)* ***b.*** *The only equality constraint to verify in this problem is the physical law of mass conservation: SV=0. Therefore, the stoichiometric matrix S is used as the matrix B in Yang et al. formulation* ***c.*** *$S_{int,In}$ was constructed to ensure two inequalities: first the uptake fluxes should be inferior to their upper bounds (UB), which is verified with the 4 first rows (corresponding to $S_{int}$, defined in Figure S4); and all fluxes should be positive, which is verified by $-I_n$ (the same as in Figure S4), that is stacked to $S_{int}$. Consequently, $S_{int,In}$ is used as the matrix A in Yang et al. formulation.*

**QP-solver equations**

We recall below the quadratic program (QP) exposed as eq. 1 in Methods 'Derivation of loss functions:

$$min( ||P_{ref}\ V - V_{ref}||^2) \tag{S10}$$

s. t. $S\ V = 0$

$P_{in}V \leq V_{in}$

$V \geq 0$

To solve this problem, a loss function with four terms was built. As mentioned in the Methods section 'Loss functions derivation', the first term is related to the fit to the reference targeted values and the three additional losses terms are related to the boundary, stoichiometric and flux positivity constraints of the metabolic network.

The first loss is simply the Mean Square Error (MSE) between predictions ($V$) and FBA-simulated or measured reference data ($V_{ref}$):

$$L_1 = \frac{1}{n_{ref}} ||P_{ref}V - V_{ref}\ ||^2 \tag{S11}$$

The second loss is linked to the network stoichiometric constraint ($S\ V = 0$), which in its normalized form (loss per constraint) is:

$$L_2 = \frac{1}{m}\ ||SV||^2 \tag{S12}$$

The third loss evaluates how well boundary constraints are respected ($P_{in}V\ \leq\ V_{in}$):

$$L_3 = \frac{1}{n_{in}} ||ReLU(\ P_{in}V - V_{in}\ )||^2 \tag{S13}$$

The last loss enforces all fluxes to be positive:

$$L_4 = \frac{1}{n} ||ReLU(-V)||^2 \tag{S14}$$

We note that when exact bounds are provided, $P_{in}V = V_{in}$, and $L_3$ becomes obsolete as the values of $V$ corresponding to $V_{in}$ are not updated by the LP/QP solvers and AMN programs.
Thus, the sum of those four terms is the loss L given be eq. 2 in Methods 'Loss functions derivation'.
Note that $L$ can also be computed as the MSE between the vectors:

$\left(P_{ref}V, \frac{||SV||}{\sqrt{m}}, \frac{||ReLU(\ P_{in}V - V_{in}\ )||}{\sqrt{n_{in}}}, \frac{||ReLU(-V)||}{\sqrt{n}}\right)$ and $\left(V_{ref}, 0, 0, 0\right)$

While the QP system can be solved by a simplex algorithm, solutions can also be approximated calculating $V$ corresponding to:

$$min\left(\frac{1}{n_{ref}}\|P_{ref}V - V_{ref}\|^2 + \frac{1}{m}\|SV\|^2 + \frac{1}{n_{in}}\|ReLU(P_{in}V - V_{in})\|^2 + \frac{1}{n}\|ReLU(-V)\|^2\right) \quad \text{(S15)}$$

The vector $V$ can thus be found solving:

$$\frac{\partial\left(\frac{1}{n_{ref}}\|P_{ref}V - V_{ref}\|^2 + \frac{1}{m}\|SV\|^2 + \frac{1}{n_{in}}\|ReLU(P_{in}V - V_{in})\|^2 + \frac{1}{n}\|ReLU(-V)\|^2\right)}{\partial V} = 0 \quad \text{(S16)}$$

As mentioned in Methods 'QP solver', $V$ satisfying eq. S15 can be found iteratively:

$$V^{(t+1)} = V^{(t)} - dt\,\nabla V \quad \text{(S17)}$$

$$V^{(0)} = P_{in}^T V_{in}$$

where $t$ is the iteration number, $dt$ the learning rate.

$\nabla V$ is computed as follow:

$$\nabla V = \frac{2}{n_{ref}} P_{ref}^T\left(P_{ref}V - V_{ref}\right) + \frac{2}{m} S^T SV + \frac{2}{n_{in}} P_{in}^T D_{in} ReLU(P_{in}V - V_{in}) - \frac{2}{n} D_V\, ReLU(-V) \quad \text{(S18)}$$

It is easy to verify that for the first term of $\nabla V$ we have:

$$\nabla V_1 = \frac{\partial \|\frac{P_{ref}V - V_{ref}}{n_{ref}}\|^2}{\partial V} = \frac{2}{n_{ref}} P_{ref}^T\left(P_{ref}V - V_{ref}\right) \quad \text{(S19)}$$

for the second term:

$$\nabla V_2 = \frac{\partial \|\frac{SV}{m}\|^2}{\partial V} = \frac{2}{m} S^T SV \quad \text{(S20)}$$

for the third term:

$$\nabla V_3 = \frac{\partial \|\frac{ReLU(P_{in}V - V_{in})}{n_{in}}\|^2}{\partial V} = \frac{2}{n_{in}} P_{in}^T D_{in} ReLU(P_{in}V - V_{in}) \quad \text{(S21)}$$

where $D_{in} = \frac{ReLU(P_{in}V - V_{in})}{ReLU(P_{in}V - V_{in})}$ using an *Hadamard* division: $\frac{A}{A} = \left(\frac{a_{ij}}{a_{ij}}\right)$ (= 0 when $a_{ij} = 0$)

and for the fourth term:

$$\nabla V_4 = \frac{\partial \|\frac{ReLU(-V)}{n}\|^2}{\partial V} = -\frac{2}{n} D_V\, ReLU(-V) \quad \text{(S22)}$$

where $D_V = \frac{ReLU(-V)}{ReLU(-V)}$ using an *Hadamard* division.

Summing eqs. S19-S22 we find:

$$\nabla V = \frac{2}{n_{ref}} P_{ref}^{T}\left(P_{ref}\, V - V_{ref}\right) + \frac{2}{m} S^{T}SV + \frac{2}{n_{in}} P_{in}^{T}\, D_{in} ReLU(P_{in}V - V_{in}) - \frac{2}{n} D_{V}\, ReLU(-V) \quad \text{(S23)}$$

In the EB case where exact medium uptake fluxes are known, the QP system is:

$$min(\, \|P_{ref}\, V - V_{ref}\|^{2}) \quad \text{(S24)}$$

s.t. $S\,V = 0$

$P_{in}V = V_{in}$

$V \geq 0$

In such an instance, reaction fluxes having a mapping in $V_{in}$ remain constant and are not updated, therefore:

$$\nabla V = \left(\frac{2}{n_{ref}} P_{ref}^{T}\left(P_{ref}\, V - V_{ref}\right) + \frac{2}{m} S^{T}SV - \frac{2}{n} D_{V}\, ReLU(-V)\right) \odot \left(1_{n} - P_{in}^{T} 1_{\{n_{in}\}}\right) \quad \text{(S25)}$$

where $\odot$ stands for *Hadamard* product (A$\odot B = a_{ij}b_{ij}$) and $1_{n}$ $\left(respectively\ 1_{\{n_{in}\}}\right)$ is a vector of dimension $n$ $(respectively\ n_{in})$ with constant coefficients equal to 1.

##### MM solvers benchmarking

As it has been described along this paper, AMNs are composed of both a mechanistic layer and a neural layer. As illustrated in Figure S6d and S6e, mechanistic solvers require more than 10,000 iterations, which brings issues to the gradient backpropagation (vanishing or exploding gradients) as well as an increased training time. Further results about mechanistic layers alone are given in the Table S1, for the 'MM' model types.

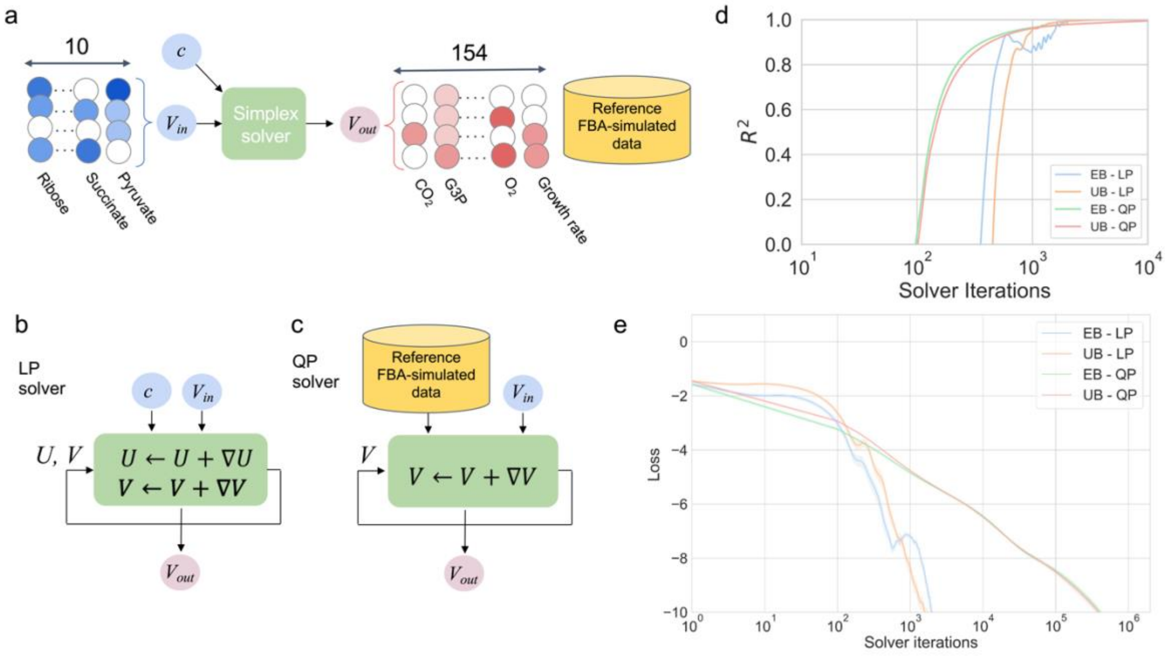


**Figure S6. MM solvers architectures and performances**

***a.*** *Schematic procedure for the Simplex solvers. From* $V_{in}$*, which is a vector describing the bounds of some uptake fluxes, the solvers reach a steady-state solution,* $V_{out}$*, optimizing the objective function c, satisfying the constraints and bounds of the network. Solutions obtained using the simplex-based method in Cobrapy*[11]*, are taken as reference data.* ***b.*** *Schematic for LP-solver architecture. This solver surrogates the simplex-based algorithm. Following Yang et al. and as further detailed in the Methods 'AMN architectures', the full flux distribution V is updated by ∇V and the metabolites shadow prices U by ∇U through products of matrices derived from the stoichiometry of the network.* ***c.*** *Schematic for QP-solver architecture. Here target reference fluxes (*$V_{ref}$*) are given to the solver. The computed fluxes are fitted to the reference targets by means of a custom loss function integrating also the input constraints along with the stoichiometric constraint of the metabolic network. The flux vector V is updated by ∇V which is the gradient minimizing the loss function (cf. Methods 'AMN architectures' for further details).* ***d.*** $R^2$ *vs Solver iteration.* ***e.*** *Loss vs. Solver iterations. QP takes 1 million iterations to reach close to zero values, whereas LP takes 10,000 iterations. In d and e, plotted is the mean and standard error (95% confidence interval) across all elements of the set of 100 simulations.*

**AMNs benchmarking varying hyperparameters**

We benchmarked hyperparameters for the neural layer of the AMN-QP model shown in Figure 2b (trained on a 1,000 *E. coli* core simulations dataset). Results are presented in Figure S7 panels b-d. The main conclusion of this search is that increasing the number of layers was inducing overfitting (displaying worst performance on validation sets than with a single layer) and that a minimal hidden dimension was 100 for better performance.

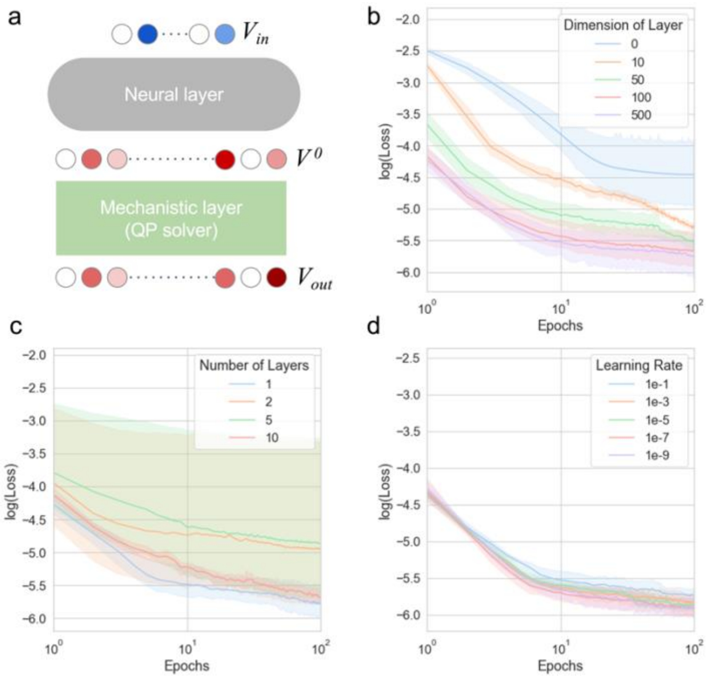


**Figure S7. Hyperparameters for AMN's neural Layer**

***a.*** *AMN architecture. In panels b-d the AMNs with QP-solver were trained on a simulated training set of 1000 samples for 100 epochs using the Adam optimizer. The metabolic model was E. coli core. Three hyperparameters were tested: dimension of the hidden layer(s), number of hidden layers and learning rate. Plotted is the mean and standard error (95% confidence interval) of the loss on test set across 5-folds cross-validation.* ***b.*** *log(Loss) vs epoch.* ***c.*** *log(Loss) vs epoch.* ***d.*** *log(Loss) vs epoch.*

From Figure S7 we note that: (i) increasing dimension of the hidden layer increases the decay of the loss function, (ii) the number of hidden layers exhibits huge variability across cross-validation folds suggesting overfitting, and (iii) no major influence was detected for the learning rate. The default architecture of the neural layer in all AMN was therefore set as one layer of dimension higher than 50 (we increased this hyperparameter with the model's size, see Table S1) and a training rate of 1e-3

#### AMNs benchmarking with independent test sets and additional metabolic models

The performances of all AMN architectures (Wt, LP, QP) are given in Table S1 using FBA simulated data on two different *E. coli* metabolic models, *E. coli* core model[4] and iML1515[2] along with iJN1463 *P. putida* model. These *E. coli* models are composed respectively of 154 reactions and 72 metabolites, and 3682 reactions and 1877 metabolites (after duplicating bidirectional reactions). The *P. putida* model is composed of 2135 reactions and 1637 metabolites (after duplicating bidirectional reactions). In all cases the default Simplex-based solver (GLPK) of Cobrapy was run to optimize growth rates for different media. Each medium was composed of metabolites found in minimal media (M9) and additional metabolites (sugars, acids) crossing the cell membrane (more details in Methods 'Generation of training sets with FBA'). For comparison purposes, Table S1 also provides results for MM architectures (no neural layer) and ANN architectures (no mechanistic layer).

**Table S1. Benchmarking MMs, ANNs and AMNs**

*(1) All SBML models describing different E. coli strains were downloaded from the BiGG database, 'Core' stands for the E. coli core model, EB (UB) stands for exact bounds (upper bounds) for medium uptake fluxes, the iML1515 model was reduced following the procedure described in Methods 'Making metabolic networks suitable for neural computations'. (2) Training set size (number of elements multiplied by number of labelled data per element) and range for the number of metabolites added to the minimal medium. (3) YES or NO if the model contains a neural layer or a mechanistic layer. (4) MM stands for Mechanistic Model, ANN stands for Artificial Neural Network (a dense neural architecture) and AMN for Artificial Metabolic Network. ANN and AMN architectures are described in Methods. Neural Layer Hyperparameters display the number of hidden layers, the size of the hidden layer, and the training rate. Mechanistic Layer Hyperparameters display the number of iterations performed by the solver. (5) Number of trainable parameters and epochs, in all cases dropout = 0.25, batch size = 5, the optimizer is Adam and the loss function is the mean squared error between predicted and reference fluxes to which for AMN loss constraints are added, see Methods 'Loss functions derivation' for additional details. (6) Regression coefficient and Loss values for training set ($R^2$), and cross-validation sets ($Q^2$) between reference growth rate and predicted growth rate. (7) Regression coefficient and Loss values for growth rates for independent test sets not seen during training. Test set sizes are 10% of training set sizes. For (6) and (7) the performance is displayed as the mean over 5 folds (or over a training set when no cross-validation scheme is performed, i.e., for the MM performances). n/a: not applicable or not computed.*

| SBML strain<br>Bound<br>(1) | Size<br>Range<br>(2) | Neural layer<br>Mechanistic layer<br>(3) | Architecture<br>Neural Layer Hyperparameters<br>Mechanistic Layer Hyperparameters<br>(4) | Nbr param.<br>Nbr epochs<br>(5) | Training $R^2$<br>Loss constraint<br>(6) | Validation set $Q^2$<br>Loss constraint<br>(6) | Test set $Q^2$<br>Loss constraint<br>(7) |
|---|---|---|---|---|---|---|---|
| Core<br>EB | 100<br>1-6 | NO<br>YES | MM_LP<br>n/a<br>$10^4$ | n/a<br>n/a | 1.00 ± 0.000<br>3.2e-9 ± 3.2e-8 | n/a<br>n/a | n/a<br>n/a |
| Core<br>UB | 100<br>1-6 | NO<br>YES | MM_LP<br>n/a<br>$10^4$ | n/a<br>n/a | 1.00 ± 0.000<br>5e-7 ± 2.8e-6 | n/a<br>n/a | n/a<br>n/a |
| Core<br>EB | 100<br>1-6 | NO<br>YES | MM_QP<br>n/a | n/a<br>n/a | 1.00 ± 0.000<br>7.8e-6 ± 6.1e-6 | n/a<br>n/a | n/a<br>n/a |

| | | | | | | | |
|---|---|---|---|---|---|---|---|
| | | | $10^6$ | | | | |
| Core<br>UB | 100<br>1-6 | NO<br>YES | MM_QP<br>n/a<br>$10^6$ | n/a<br>n/a | 1.00 ± 0.000<br>7.1e-6 ± 5.7e-6 | n/a<br>n/a | n/a<br>n/a |
| Core<br>EB | 1540<br>1-6 | YES<br>NO | ANN<br>1, 50, 1e-3<br>n/a | 8904<br>500 | 0.83 ± 4.8e-2<br>1.8e-1 ± 2.2e-3 | 0.66 ± 1.4e-1<br>2.6e-1 ± 2.1e-2 | n/a<br>n/a |
| Core<br>UB | 1540<br>1-6 | YES<br>NO | ANN<br>1, 50, 1e-3<br>n/a | 8904<br>500 | 0.82 ± 4.8e-2<br>1.1e-1 ± 8.2e-3 | 0.39 ± 2.5e-1<br>2.1e-1 ± 7.4e-2 | n/a<br>n/a |
| Core<br>EB | 154 000<br>1-6 | YES<br>NO | ANN<br>1, 50, 1e-3<br>n/a | 8904<br>100 | 0.91 ± 1.5e-2<br>2.3e-1 ± 2.3e-2 | 0.98 ± 1.1e-2<br>1.0e-2 ± 1.1e-2 | n/a<br>n/a |
| Core<br>UB | 154 000<br>1-6 | YES<br>NO | ANN<br>1, 50, 1e-3<br>n/a | 8904<br>100 | 0.86 ± 2.1e-2<br>1.8 ± 2.0 | 0.94 ± 3.9e-2<br>4.4e-3 ± 1.0e-3 | n/a<br>n/a |
| Core<br>EB | 1000<br>1-6 | YES<br>YES | AMN_LP<br>1, 50, 1e-3<br>4 | 17 808<br>500 | 0.98 ± 7.9e-3<br>2.8e-3 ± 0.6e-3 | 0.98 ± 7.4e-4<br>2.8e-3 ± 0.5e-3 | 0.98<br>3.0e-3 |
| Core<br>UB | 1000<br>1-6 | YES<br>YES | AMN_LP<br>1, 50, 1e-3<br>4 | 25 152<br>500 | 0.98 ± 9.7e-3<br>2.5e-3 ± 0.4e-3 | 0.97 ± 1.0e-2<br>2.5e-3 ± 0.4e-3 | 0.99<br>3.1e-3 |
| Core<br>EB | 1000<br>1-6 | YES<br>YES | AMN_QP<br>1, 50, 1e-3<br>4 | 8904<br>500 | 0.99 ± 4.2e-3<br>2.3e-3 ± 0.5e-3 | 0.99 ± 4.7e-3<br>2.3e-3 ± 0.5e-3 | 0.98<br>3.0e-3 |
| Core<br>UB | 1000<br>1-6 | YES<br>YES | AMN_QP<br>1, 50, 1e-3<br>4 | 8904<br>500 | 0.97 ± 9.9e-3<br>2.5e-3 ± 0.6e-3 | 0.97 ± 1.3e-2<br>2.5e-3 ± 0.6e-3 | 0.97<br>2.0e-3 |
| Core<br>UB | 1000<br>1-6 | YES<br>YES | AMN_Wt<br>1, 50, 1e-3<br>4 | 13 622<br>500 | 0.99 ± 1.3e-3<br>0.9e-3 ± 0.000 | 0.99 ± 2.2e-3<br>0.9e-3 ± 0.000 | 1.0<br>0.000 |
| iML1515<br>UB | 11000<br>1-6 | YES<br>NO | ANN<br>1, 500, 1e-3<br>n/a | 295 050<br>100 | 0.88 ± 4.3e-2<br>2.2 ± 0.8 | 0.76 ± 1.0e-1<br>4.7 ± 4.2 | n/a<br>n/a |
| iML1515<br>UB | 550 000<br>1-6 | YES<br>NO | ANN<br>1, 500, 1e-3<br>n/a | 295 050<br>100 | 0.98 ± 3.3-2<br>4.0e-4 ± 3.0e-4 | 0.67 ± 3.5e-1<br>3.1e-3 ± 4.5e-3 | n/a<br>n/a |
| iML1515<br>UB | 11000<br>1-4 | YES<br>YES | AMN_LP<br>1, 250, 1e-3<br>4 | 839 266<br>100 | 1.0 ± 1.0e-3<br>0.000 ± 0.000 | 1.0 ± 1.0e-3<br>0.000 ± 0.000 | 1.0<br>0.000 |

| | | | | | | | |
|---|---|---|---|---|---|---|---|
| iML1515<br>UB | 11000<br>1-4 | YES<br>YES | AMN_QP<br>1, 500, 1e-3<br>4 | 295 050<br>100 | 1.0 ± 1.4e-3<br>0.000 ± 0.000 | 1.0 ± 1.4e-3<br>0.000 ± 0.000 | 1.0<br>0.000 |
| iML1515<br>UB | 11000<br>1-4 | YES<br>YES | AMN_Wt<br>1, 500, 1e-3<br>4 | 634 238<br>100 | 1.0 ± 0.1e-3<br>0.000 ± 0.000 | 0.99 ± 0.4e-3<br>0.000 ± 0.000 | 1.0<br>0.000 |
| iJN1463<br>UB | 4860<br>1-1 | YES<br>YES | AMN_QP<br>1, 500, 1e-3<br>4 | 1 168 135<br>500 | 0.99 ± 2e-3<br>0.000 ± 0.000 | 0.99 ± 2e-3<br>0.000 ± 0.000 | 0.99<br>0.000 |

The MM architectures show good performances both in terms of growth rate computation and loss on constraints. There is no learning process involved with MMs, therefore no reason to compute results for validation and test sets. The ANN architectures (cf. Methods 'ANN architecture' for further details) exhibit poor performances and have small predictive capacities (high Loss) for cross validation sets of sizes in the range of those of AMN (~1000 reference data in training), for that reason performances for test sets were not assessed. We also note that losses remain higher with ANNs even for training set sizes 1000 times larger than those used with AMNs. All AMN architectures exhibit excellent regression coefficients and losses for training sets, validation sets and test sets, and this for both models *E. coli* core[4], iML1515[2] and iJN1463[12].

**AMNs benchmarking with gene knockouts and multiple measured fluxes**

To assess the performance of AMNs on datasets where more than one flux is measured, we extracted a dataset from Rijsewijk *et al.*[13] which consists of 128 experiments, each containing 31 measured fluxes.

The dataset was composed of 2 media compositions (glucose or galactose as carbon source), for 64 regulator gene KOs mutants ($G_{KO}$). These regulator genes were found on RegulonDB[14] and their corresponding regulated metabolic reactions encoded in iML1515 were compiled. Each regulator was found to have at least one regulated reaction in iML1515. The final training set to use with AMNs was composed of 2 inputs: 1 binary vector of size 2 for media compositions ($C_{med}$) and 1 binary vector of size 64 for gene KOs ($G_{KO}$). Unlike for the *E. coli* KO dataset used in Figure 4 in the main paper, we did not add a term to the custom loss since the effect of deleting the regulation of a reaction is *a priori* unknown (at least quantitatively, in terms of effect on the fluxes distribution).

Overall, the performance is satisfactory for most fluxes, but 7 fluxes (empty slots in Figure S8b) have a $Q^2$ close to zero or negative. However, these low-performance predictions do not impact the variance-weighted average $Q^2$ (red line, 0.91) because the corresponding measured fluxes have low variance, thus, there are limited statistical patterns for the model to learn on these fluxes (we recall that the variance weighted-average consists in a weighted average of all 31 fluxes' $Q^2$s, with a weight applied on each flux's $Q^2$ corresponding to the variance found in the flux's measure)s.

Learning on many fluxes is more challenging for the AMNs than on a single flux, but it still seems to make accurate predictions with this dataset for the majority of fluxes.

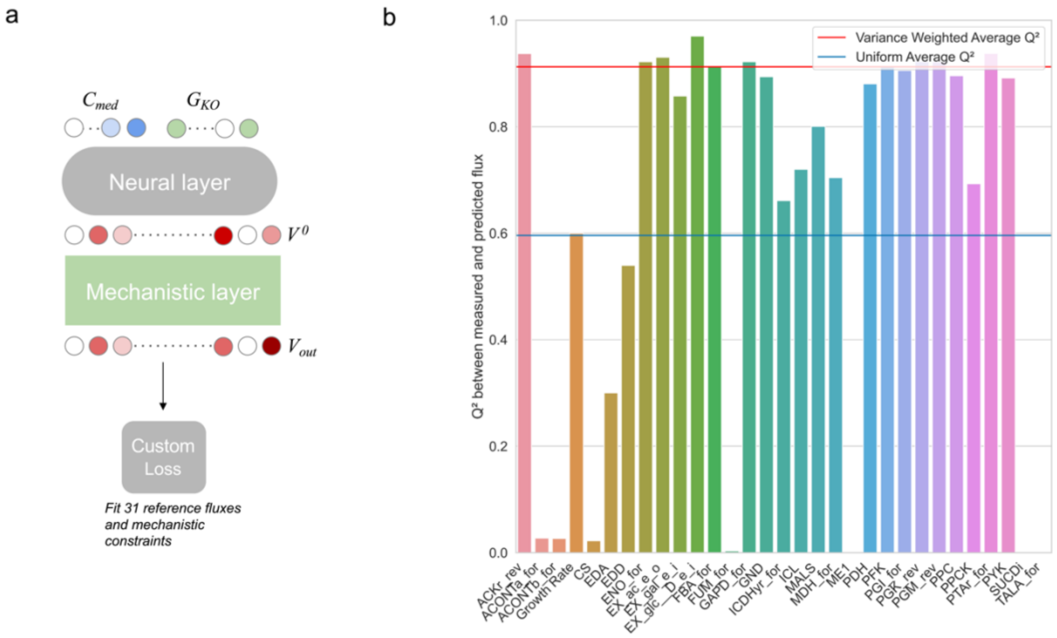


#### Figure S8. AMN performance on multiple fluxes dataset

***a.*** *AMN architecture compatible with the multiple fluxes dataset from Rijsewijk et al. The mechanistic layer and custom loss were derived from the E. coli model iML1515. The medium composition binary vector for this dataset is of size 2, for glucose or galactose as the carbon source. The other input,* $G_{KO}$*, a binary vector as well, of size 64, is describing which genes are knocked out. The remaining of the architecture is similar to what is shown in Figure 1c, with 2 hidden layers of size 400 as the neural layer, QP as the mechanistic layer, and a custom loss fitting simultaneously 31 reference fluxes (instead of only the growth rate for most results shown in the study). The model was trained for 100 epochs with the Adam optimizer with a learning rate of 1.0e-3.* ***b.*** *Performance chart of the AMN, displayed as* $Q^2$ *computed on aggregated validation sets from a 10-fold cross-validation. The mean predictions over 3 repeats of the 10-fold cross-validation were compiled as final prediction values. The* $Q^2$ *is displayed for each flux individually with the bars of the chart, and we also show the variance weighted (red line, 0.91) and uniform (blue line, 0.60) averages of* $Q^2$*s of all fluxes.*

**AMN-Reservoir prediction performance**

Figure 5 in the main paper displays the performance of classical FBA with $V_{in}$ extracted from the AMN-Reservoir, after training on the whole dataset. Another possibility, is to use the AMN-Reservoir in a more predictive manner, obtaining $V_{in}$ during predictions on the validation sets of a cross-validation. We show in Figure S9 below the performance of FBA when using such predicted $V_{in}$ from unseen data, as the regression performance on the 110 *E. coli* growth rates dataset. We note the regression coefficient we obtained is similar to those obtained when training AMNs directly on experimental data (Figure 3, main paper).

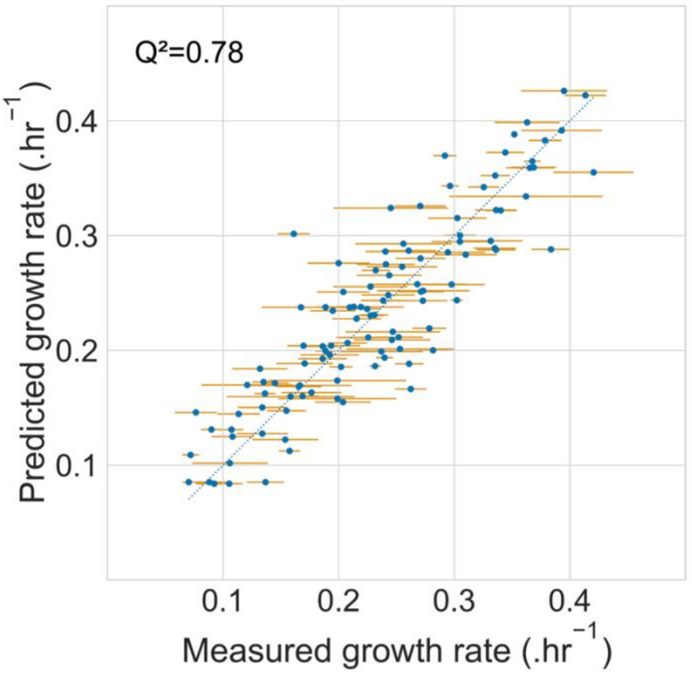


**Figure S9. Performances of AMN-Reservoir using predicted $V_{in}$ as input to FBA**

*The dataset used to train the AMN-Reservoir was the 110 E. coli growth rates used for Figure 3 and Figure 5 panels c and d. The measured growth rates are plotted as the mean and standard deviation over technical replicates (cf. Methods). The hyperparameters and the pre-trained AMN-Reservoir were the same as for Figure 5 panel c. A 10-fold cross-validation was performed (instead of a training and prediction on whole dataset as in Figure 5c), and validation sets predictions were used to extract $V_{in}$ then use it as input for FBA. FBA results are shown here as the predicted growth rate.*

##### ANN training set sizes

To compare the performances of an ANN 'black box' model with AMNs, we trained a simple dense ANN model and an AMN-QP model on training sets of increasing sizes. The training sets were generated using *E. coli* core[4] as in Methods 'Generation of training sets with FBA'. We recall (see Methods 'ANN architecture') that to assess losses for an ANN model (which does not have any mechanistic layer), each entry of the training sets contains all flux values; this enables one to compute the losses given in Methods 'Loss function derivation'. Consequently, with ANN for each element in the training set we provided as labeled data all reaction fluxes (154 for *E. coli* core[4]), while with AMN-QP we provided as labeled data only the flux of the biomass reaction.

To enable comparison between AMN and ANN, in Figure S10 given below the training set size is the number of labeled data provided. We find that the results obtained for AMN-QP are consistent with those presented in Table S1: training set size of 1000 yields a $Q^2$ above 0.95 while the loss remains below 0.003. We also observe that the ANN architecture requires training set sizes several orders of magnitude larger to reach losses that are still above those obtained with AMN-QP: training set sizes of more than 500,000 are needed to obtain a loss below 0.01, while $Q^2$ above 0.9 are reached for training set sizes above 50,000. Finally, it is worth noticing that while ANN can be trained on simulated data as in Figure S10 they cannot directly be used with experimental data as it is practically not possible to measure all the reaction fluxes of a strain grown with different media compositions.

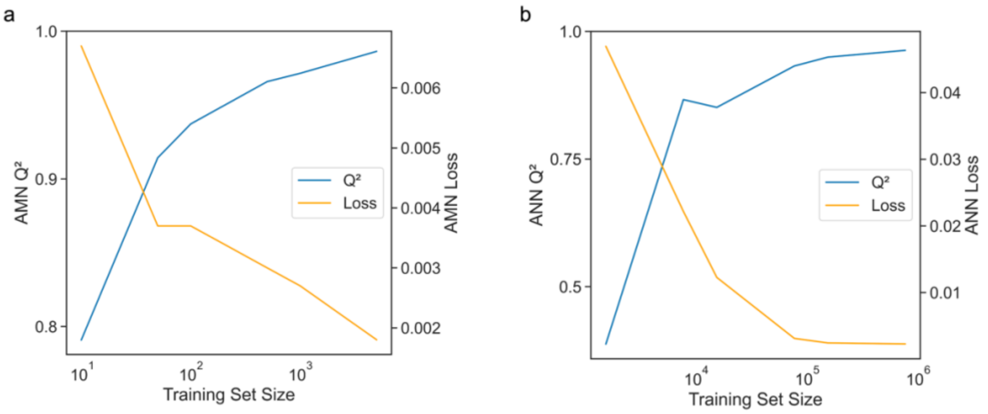


**Figure S10. Loss and regression coefficient for training sets of increasing sizes**
*In both cases training sets were generated for the E. coli core model using the procedure described in Methods 'Generation of training sets with FBA'. AMN and ANN were trained for different medium metabolites uptake rates as inputs and, as reference (labeled) data, the biomass reaction flux for AMN and all fluxes for ANN. In both cases Q² is the regression coefficient between the reference and the predicted biomass reaction fluxes during 5-fold cross-validation. Loss on constraints were computed as described in Methods 'Loss functions derivation' on 5-fold cross-validation sets.* ***a.*** *AMN. The model architecture is the one shown in Figure 3 with a QP-solver for the mechanistic layer. The neural layer is composed of an input layer of size 20 (all uptake fluxes of E. coli core), a hidden layer of size 50 and an output layer of size 154 (all reactions of E. coli core), the learning rate was set to 1.0e-3 and Adam was used as the optimizer.* ***b.*** *ANN. The ANN model has the same architecture as the neural layer of the AMN and no mechanistic layer. Raw data for this figure can be found in the amn_release GitHub repository (Result folder).*

#### Experimental workflow

Details on the experimental protocol can be found in the Methods 'Culture conditions' and 'Growth rate determination'. Figure S11 gives a visual overview of the workflow to generate the dataset showcased in Figures 3 & 5 (main paper).

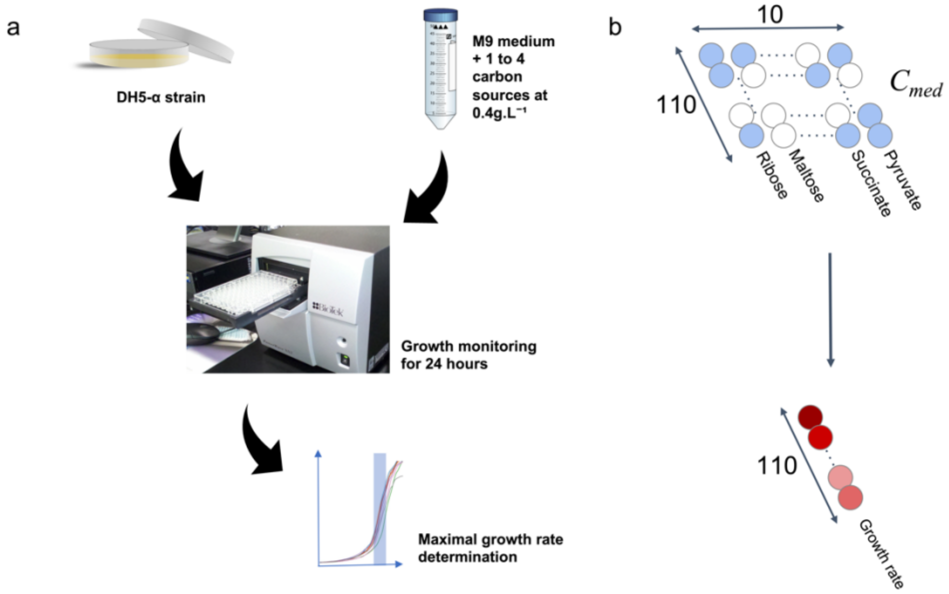


**Figure S11. Experimental workflow pipeline**
***a.*** *The DH5-alpha strain of E. coli was cultured in M9 medium with different carbon source combinations (1 to 4 carbon sources simultaneously added, all at 0.4g.L*$^{-1}$*). The optical density at 600nm (OD600) was monitored for 24 hours in a plate reader; reading 96-well plates each containing 10 media compositions, each in 8 replicates (remaining space was used as blanks, for the edges of the plate that show high evaporation). After data acquisition, the maximal growth rate was computed (cf. Methods 'Growth rate determination').* ***b.*** *The experimental workflow enables the generation of 110 data points each composed of* $C_{med}$ *as the independent variables and growth rates as dependent variables.* $C_{med}$ *is describing each medium carbon source composition as zeros - when absent - and ones - when present – yielding in the end a binary vector of length 10.*

**Terminology**

We provide below the list of all notations used in all equations and figures of our main manuscript and Supplementary Information.

**Table S2. Vectors and matrices notations used in figures and equations**

| **Notation** | **Description (units)** |
|---|---|
| $V_{in}$ | **Vector** of exact or upper bounds for uptake fluxes. See Figure 1a,c,d (mmol.gDW$^{-1}$.h$^{-1}$) |
| $V_{out}$ | **Vector** of steady-state fluxes values predicted by a model, either fully mechanistic or AMN. See Figure 1. (mmol.gDW$^{-1}$.h$^{-1}$ and .h$^{-1}$) |
| $V_{ref}$ | **Vector** of reference fluxes, either FBA-simulated or measured. In all results except Figure S8 (31 fluxes) and for the ANNs (all fluxes), it contains only the growth rate. (mmol.gDW$^{-1}$.h$^{-1}$ and .h$^{-1}$) |
| $V^0$ | **Vector** of fluxes values before passing through the mechanistic model. Referred to as initial guess for the flux distribution. See Figure 1c,d. (mmol.gDW$^{-1}$.h$^{-1}$ and .h$^{-1}$) |
| $C_{med}$ | **Vector** describing medium composition. See Figure 1c,d. (no unit) |
| V | **Vector** of reaction fluxes. Generic name. (mmol.gDW$^{-1}$.h$^{-1}$ and .h$^{-1}$) |
| M | **Vector** of metabolites production rates, used for Wt. (mmol.gDW$^{-1}$.h$^{-1}$) |
| U | **Vector**. Dual variable of V when considering FBA's constrained linear problem. Also called metabolites' shadow prices. |
| $R_{KO}$ | **Vector** of reactions that are KO. See Figure 4a. 0 if reaction is inactivated by the gene KO, 1 otherwise. (no unit) |
| $c_{FBA}$ | **Vector** of reactions that are hypothesized to be maximized by the cell. In this work, always set to the biomass production reaction (*i.e.,* the growth rate). Used for LP method. (mmol.gDW$^{-1}$.h$^{-1}$) |
| S | **Matrix** of stoichiometric coefficients given by the GEM. Its dimension is *m* (number of metabolites) × *n* (number of reactions). (no unit) |
| $W_r$ | **Matrix** of weight representing consensual flux branching ratios. (no unit) |
| $P_{in}$ | **Matrix** of mapping V into the fluxes of $V_{in}$. (no unit) |
| $P_{out}$ | **Matrix** of mapping V into the fluxes of $V_{out}$. (no unit) |
| $P_{ref}$ | **Matrix** of mapping V into the fluxes of $V_{ref}$. (no unit) |
| $P_{Ko}$ | **Matrix** of mapping from gene KO to inactivated reactions. (no unit) |
| $P_{v\rightarrow m}$ | **Matrix** of mapping from reactions to metabolites. (no unit) |
| $P_{m\rightarrow v}$ | **Matrix** of mapping from metabolites to reactions. (no unit) |
| $S_{int}$ | **Matrix** of stoichiometric coefficients where uptake reactions have been zeroed out. (no unit) |

In our study $V_{in}$ are "uptake fluxes" also named "uptake reactions" the fluxes and corresponding reactions that introduce matter into the model, *i.e*. reactions that have no reactants and their product is a metabolite in the 'medium' compartment of the metabolic model. These reactions are also called "exchange reactions" in many studies, and this subsection aims to clarify the use of "uptake flux" in this study. The term "uptake" was preferred to "exchange" for two main reasons: (i) for simplicity to readers that are not familiar with metabolic models and (ii) for the better biological sense of "uptake reactions” (designating an organism uptaking nutrients from its environment), even if these reactions introducing matter into the model are fully virtual reactions without any biological or physical sense.

Importantly, ‘uptake reactions’ are not referring to the membrane-crossing reactions, which are always left with default bounds in this study. In practice, when one makes *ab initio* predictions with classical FBA, one sets a non-zero upper bound for a reaction introducing matter in the system, to simulate the presence of a given metabolite in the medium. But, in most cases, this reaction flux optimized value will be equal to the membrane-crossing flux value, since one metabolite, in most cases, can only go to this reaction once it has been introduced in the medium compartment of the model. The only exception is in some models where the metabolites can interact in the medium compartment, or when several transport reactions are available, which is rare and not the case in the scope of the models used in this study.
